# Arachidonic acid does not grease the exocytotic SNARE machinery

**DOI:** 10.1101/2023.02.06.527244

**Authors:** Dirk Fasshauer

## Abstract

It has been proposed that arachidonic acid stimulates neurite outgrowth by specifically activating syntaxin 3, a SNARE proteins that mediates the fusion of transport vesicles with the plasma membrane and is thus instrumental for the insertion of new membrane needed for cell growth and proliferation. Considering the important role of arachidonic acid and other polyunsaturated fatty acids for human health, these findings would have wide implications for everyday life. Here, I have report that the effect of arachidonic acid is caused by a nonspecific, detergent-like action on SNARE proteins *in vitro*. The action requires unphysiologically high concentrations, is mimicked by several detergents or detergent-like substances forming micelles regardless of whether arachidonic acid is present or not, and is thus highly unlikely to be of any physiological significance for SNARE function.

## Introduction

Zipper-like assembly of <u>N</u>-ethylmaleimide-<u>s</u>ensitive factor <u>a</u>ttachment <u>re</u>ceptors (SNARE) proteins into a membrane-bridging complex is thought to drive membrane fusion reactions in the secretory pathway. SNARE-mediated fusion is controlled by a plethora of mechanisms (for reviews see (Rizo, 2022)(Jahn and Fasshauer, 2012)). The influence of membrane lipids and their derivatives has so far thought to be confined to determining the physical properties of the fusing bilayers such as facilitating the formation of highly curved intermediates. This includes free fatty acids whose concentration in membranes are thought to be low and which support negative curvature (McMahon and Gallop, 2005; Zimmerberg and Chernomordik, 2005). Arachidonic acid (AA), an essential fatty acid, is released from phospholipids by the action of phospholipase A2. Due to its four *cis* double bonds, AA favors disordered states and increases the fluidity of membranes. AA is also known to be a precursor for eicosanoid hormones that trigger specific signaling cascades (for review (Brash, 2001)).

It was shown that externally added AA reproduces the stimulating effect of nerve growth factor (NGF) on neurite outgrowth in PC12 cells (Darios and Davletov, 2006). By specific knock-downs they demonstrate that membrane expansion is mediated by the SNARE protein syntaxin 3. This protein belongs to a group of syntaxins known to be localized to the plasma membrane and to participate in a variety of different secretory processes. The authors then report that *in vitro* syntaxin 3 only binds to its partner SNARE SNAP-25 when AA is present (Darios and Davletov, 2006).

How do the authors envision the activation of syntaxin 3? Plasma membrane syntaxins can adopt two different conformations (Rizo and Sudhof, 2002). In the “closed” conformation, the regulatory N-terminal domain binds onto the C-terminal SNARE complex forming region, the SNARE motif, whereas in the “open” form the two domains are not in contact with each other. Importantly, syntaxins in the closed conformation cannot interact with their partner SNAREs. Therefore, Darios and Davletov suggest that AA causes syntaxin 3 to switch from a closed to an open conformation.

Syntaxin 3 appears also to be activated by detergents (Bajohrs et al., 2005). Surface active agents are known to affect protein conformations and the strength and kinetics of protein-protein interactions (Chandler, 2005). Thus, Davletov and colleagues seemingly have discovered two different agents that can ‘open’ syntaxins: detergents and arachidonic acid. However, arachidonic acid is amphipathic and shares chemical properties with anionic detergents (Brash, 2001). To exclude that the activating effect is caused by the surfactant properties of AA rather than by a specific interaction, the authors argue that detergents needed to be employed at millimolar concentrations (≈ 1.5 mM for Triton X-100, ≈ 27 mM for octylglucoside) (Bajohrs et al., 2005) while the maximal effect of arachidonic acid was already observed at concentrations of ≈ 100 μM (Darios and Davletov, 2006), arguing for a specific effect of AA on syntaxin 3.

Detergents are soluble in water, but when a certain threshold concentration is reached (termed critical micellar concentration, CMC), they form micelles. The surfactant properties of detergents only become apparent at concentrations above the CMC. Each detergent possesses its own unique CMC. Whereas the CMCs of Triton X-100 and octylglucoside are around 0.23 mM and 20 mM, respectively, the CMC of AA is known to be in the range of only 60 μM (Serth et al., 1991). Thus, the sigmoidal concentration dependence of AA-induced syntaxin activation (Darios and Davletov, 2006) could very likely correspond to the formation of micelles. Similarly, other polyunsaturated fatty acids shown to activate syntaxin 3 (Darios and Davletov, 2006)were used above their respective CMCs. In contrast, fatty acids used as negative control (Darios and Davletov, 2006) were all employed at concentrations below their CMC or at concentrations above their solubility level (Serth et al., 1991; Small, 1986). Thus it is conceivable that the activating effect is solely due to the presence of micelles and not related to any specific effect of AA.

## Results and Discussion

I have therefore re-investigated whether SNARE assembly is promoted by arachidonic acid and whether this promotion is due to a specific effect or due to the presence of surfactant micelles irrespective of their chemical composition. First, I found that syntaxin 3, synaptobrevin 2, and SNAP-25 readily form SNARE complexes in the absence of detergent (Fig. 1a). Similarly, no detergent was necessary for productive binary interaction of syntaxin 3 and SNAP-25 as visualized by non-denaturing polyacrylamide gel electrophoresis (Fig. 1b). Thus, formation of SNARE complexes does not require the presence of surfactants, in agreement with a previous observations of our group (Fasshauer et al., 1999). However, I noticed a slight acceleration of complex formation in the presence of 200 μM AA (Fig. 1a). To measure the rate of SNARE assembly more precisely, I used a fluorescence-based assay. Mixing of syntaxin 3 (aa 1-260) with SNAP-25 labelled at position 130 with Texas Red in the absence of detergent led to an increase in fluorescence anisotropy, denoting complex formation (Fig. 1c). Again, I observed a somewhat faster reaction in the presence of 200 μM AA (Fig. 1c). This effect, however, was also seen in the presence of nonionic detergents like Triton X-100 (data not shown) or n-Dodecyl β-D-maltoside (DDM) (Fig. 1c). Importantly, the formation of mixed micelles consisting of DDM and AA, or of CHAPS and AA, did not lead to an additional acceleration (Fig. 2a, and data not shown). Furthermore, a comparable acceleration was observed in the presence of AA or DDM when a truncated version of syntaxin 3 (aa 183-261) was used, which lacks the N-terminal regulatory domain and only contains the SNARE motif (Fig. 2b). Moreover, DDM and AA also accelerated the interaction between syntaxin 4 and SNAP-25 (Fig. 2c), i.e. the effect is not restricted to syntaxin 3. Again, acceleration was not dependent on the presence of the N-terminal domain (Fig. 2d). Very similar results were obtained when CD-spectroscopy was used as readout, which monitors the increase in α-helical content upon formation of binary and ternary SNARE complexes (Figs. 3 and 4). Finally, I investigated whether a mono-unsaturated fatty acid, oleic acid, increases the rate of SNARE complex formation when applied at concentrations above its CMC. Using both fluorescence anisotropy and CD-spectroscopy, an acceleration of assembly was observed that was similar to that observed in the presence of AA (Fig. 4b).

**Fig. 1:**
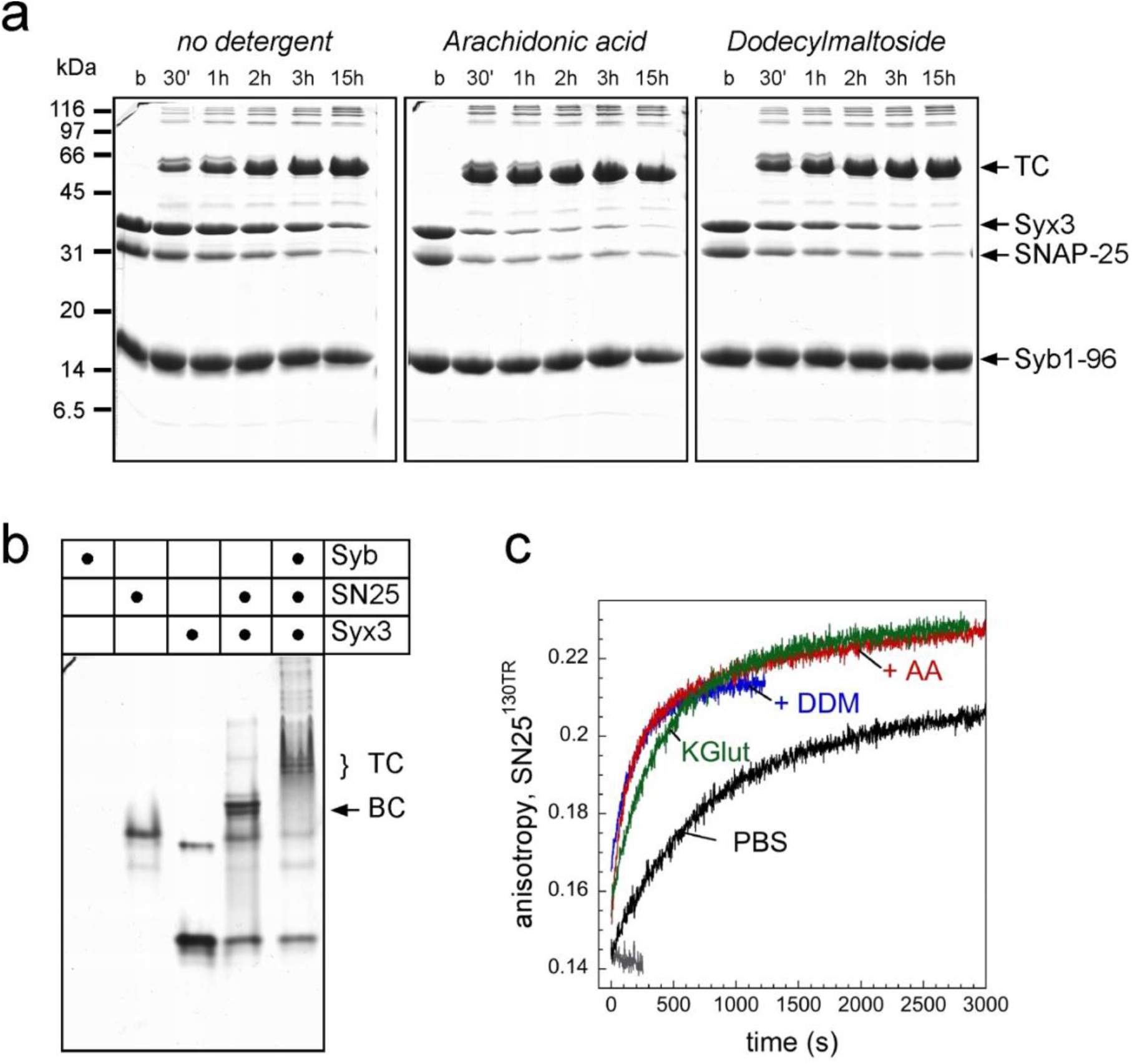
Syntaxin 3, SNAP-25, and synaptobrevin 2 readily form a SNARE complex in the absence of detergents. **a**, To monitor the formation of an SDS-resistant ternary complex (TC) approximately 20 μM syntaxin 3 (aa 1-260, Syx3), 20 μM SNAP-25, and 40 μM synaptobrevin (Syb1-96) were mixed and at each time point aliquots were added to SDS-sample buffer to stop further assembly. The reactions were carried out in PBS buffer (20 mM phosphate buffer, pH 7.4, 150 mM NaCl, 1 mM DTT) without detergent (left panel), 200 μM arachidonic acid (middle panel) or 200 μM n-Dodecyl β-D-maltoside (DDM). **b**, Syntaxin 3 readily formed a binary complex (BC) with SNAP-25 and a ternary complex (TC) with SNAP-25 and synaptobrevin monitored by non-denaturing electrophoresis. Approximately stoichiometric amounts of the proteins were mixed for one hour at room temperature before loading onto the gel. Note that due to its isoelectric point, monomeric synaptobrevin did not run into the native gel. **c**, Binary complex formation of SNAP-25 with syntaxin 3 (aa 1-260, ≈ 12 μM) was followed by a change in fluorescence anisotropy of ≈ 125 nM SNAP-25 labelled at the single cysteine 130 with Texas Red (SNAP-25^130TR^). Assembly was first carried out in PBS. In the presence of 200 μM arachidonic acid (AA) or of 200 μM DDM, a non-ionic detergent with low CMC (≈ 0.17 mM), an acceleration of assembly was observed. A similar acceleration was monitored in the presence of 0.33 % (v/v) Triton X-100 (data not shown). Note, that the detergent-free glutamate containing buffer (KGlut; 140 mM potassium glutamate, 20 mM potassium acetate, 20 mM Hepes, 2 mM EDTA, pH 7.2) also allowed for more rapid assembly (Siddiqui et al., 2007).

**Fig. 2:**
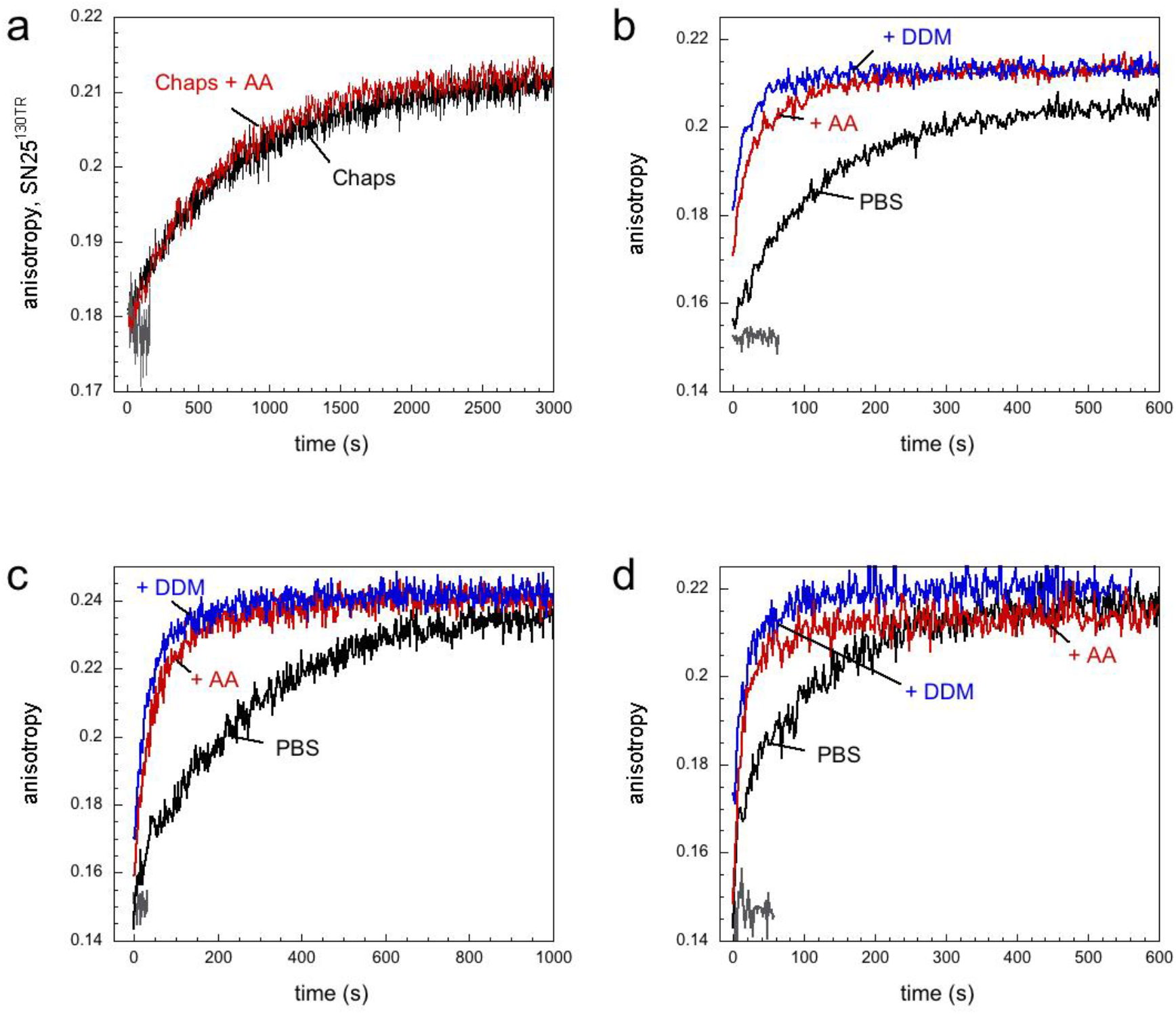
Arachidonic acid and n-Dodecyl β-D-maltoside accelerate assembly of the SNARE motifs of syntaxin 3 and syntaxin 4. **a**, In the presence of 0.67 % (w/v) Chaps in PBS buffer no acceleration of assembly of 125 nM SNAP-25^130TR^ and 12 μM syntaxin 3 was observed. Formation of mixed micelles by addition of 200 nM arachidonic acid did not accelerate assembly. **b**, Addition of AA or DDM led to an acceleration of the interaction of the SNARE motif of syntaxin 3 (aa 183-261, ≈ 3 μM) with SNAP-25^130TR^. Note that compared to the entire cytoplasmic domain of syntaxin 3 (Fig. 1c), which has to ‘open’ first to engage in SNARE interactions, its SNARE motif assembles much more quickly. **c**, Syntaxin 4 (aa 1-273, ≈ 10 μM) readily formed a complex with SNAP-25^130TR^. Assembly was accelerated in the presence of 200 μM AA or 200 μM DDM. **d**, Interaction of the SNARE motif of syntaxin 4 (aa 192-279, ≈ 3 μM) with SNAP-25^130TR^ was accelerated in the presence of 200 μM AA or 200 μM DDM.

**Fig. 3:**
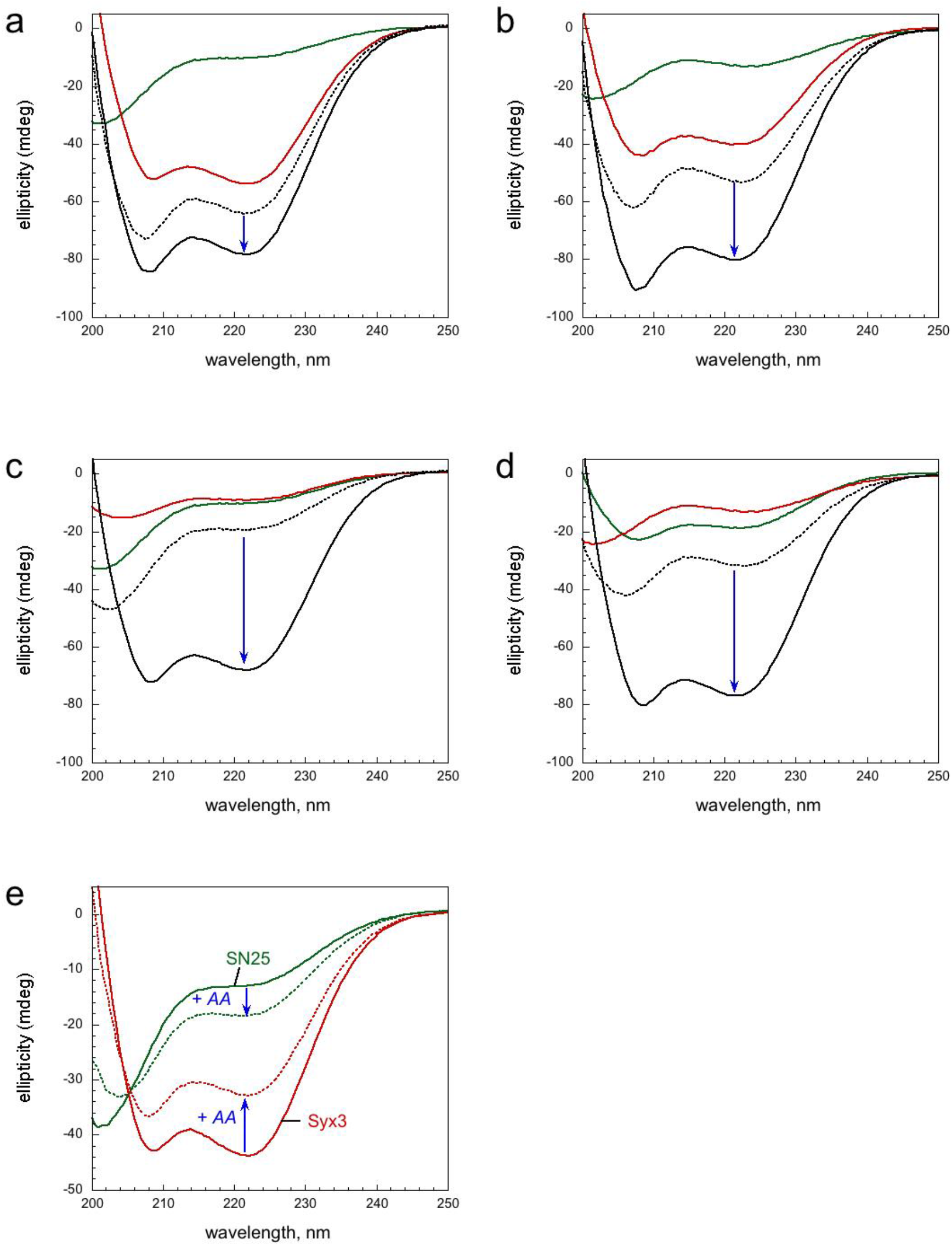
Similar structural changes observed by CD spectroscopy upon assembly of syntaxin 3 and SNAP-25 in the presence or absence of arachidonic acid. **a,** A structural change was observed when 15 μM syntaxin 3 were mixed with 12 μM SNAP-25 in PBS buffer. Green spectrum, SNAP-25; red spectrum, syntaxin 3 (aa 1-260), black spectrum, observed spectrum after 24 h incubation of SNAP-25 and syntaxin 3; dotted spectrum, theoretical sum of the spectra of SNAP-25 and syntaxin 3; the structural change is indicated by an arrow. **b,** A similar structural changes was observed in the presence of 200 μM arachidonic acid. Note, that the final spectra upon binary complex formation in the absence (a) and in the presence (b) of arachidonic acid are almost identical. The structural change observed in the presence of arachidonic acid is larger. However, this is due to the reduced α-helical content in syntaxin 3 in the presence of arachidonic acid (see e). **c + d,** The structural change was more pronounced when only the SNARE motif of syntaxin 3 (aa 183-261, 15 μM) was mixed with SNAP-25 (12 μM) in the absence (c) or presence of (d) 200 μM arachidonic acid. **e,** Arachidonic acid induced structural changes in syntaxin 3 and in SNAP-25. The spectra were first recorded in the absence of arachidonic acid and again recorded after addition of 200 μM arachidonic acid into the same cuvette. The solvent of archidonic acid, ethanol, had no significant effect on the spectra. Furthermore, oleic acid induced similar structural changes as arachidonic acid in all individual SNARE proteins (data not shown).

**Fig. 4:**
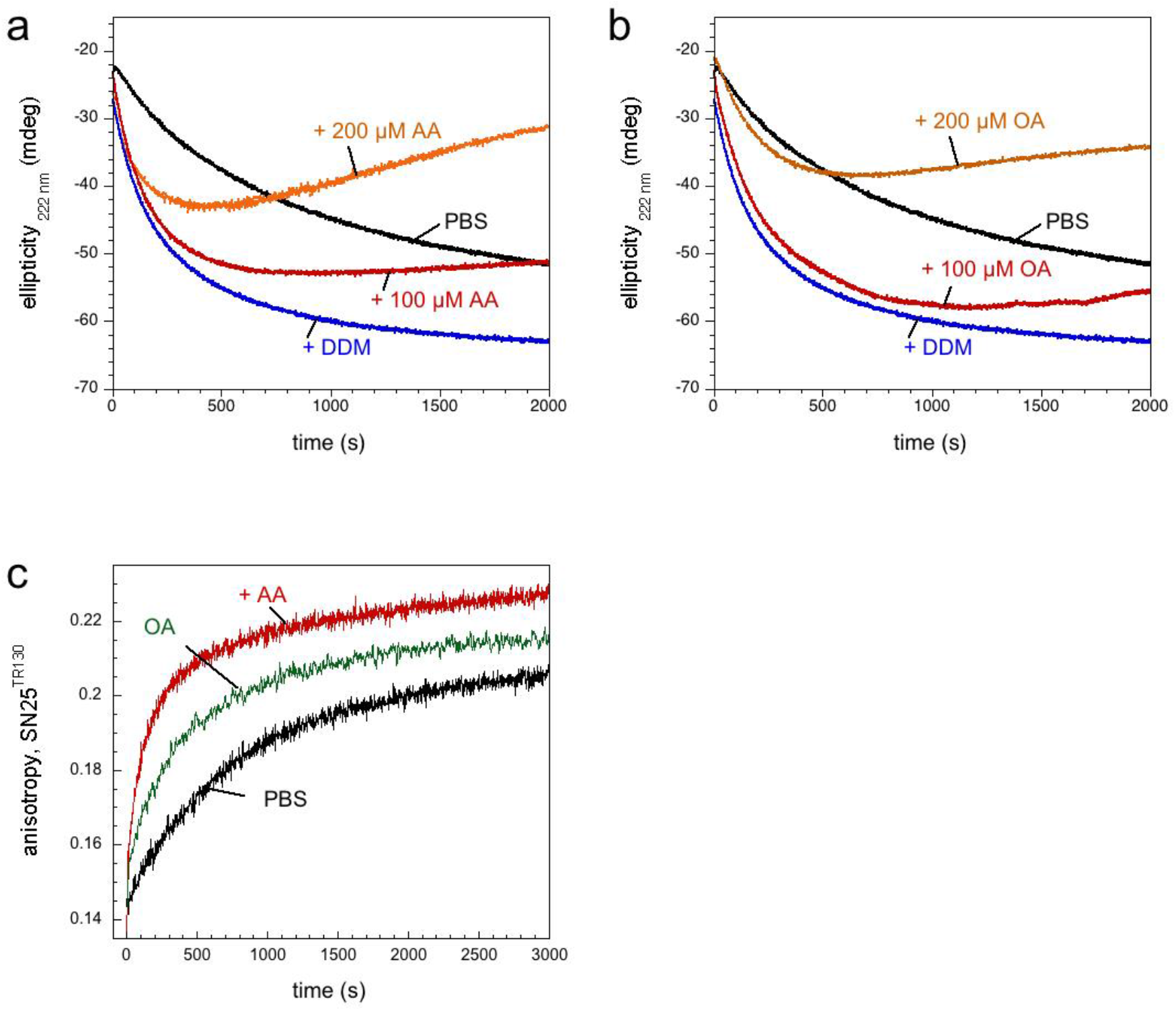
Oleic acid, a monounsaturated fatty acid, exhibits a comparable accelerating effect on SNARE assembly as arachidonic acid. **a,** The increase in α-helical content upon mixing of 15 μM of the SNARE motif of syntaxin 3 (aa 183-261) and 12 μM SNAP-25 in PBS buffer was monitored over time at 222 nm by CD spectrosopy. A clear acceleration of the reaction was observed in the presence of 200 μM DDM. Addition of 100 μM or 200 μM arachidonic acid also speed up assembly. However, the more arachidonic acid was added the more pronounced was a reversal of the change of the CD signal after several minutes of incubation, indicative for a loss of α-helical structure. After the reaction, a cloudy precipitation was observed in the cuvette, revealing that the protein partially precipitated. **b,** Oleic acid led to comparable acceleration of assembly of the SNARE motif of syntaxin 3 and 12 μM SNAP-25. In addition, similar protein precipitation as for arachidonic acid was observed. **c,** Addition of 100 μM oleic acid slightly increased the speed of interaction between SNAP-25^130TR^ and the entire cytoplasmic region of syntaxin 3 (aa 1-260, ≈ 12 μM). Higher concentrations of oleic acid appeared to quicken the interaction but also led to further protein precipitation (data not shown).

Taken together, my data show that (i) SNARE complex formation proceeds in the absence of detergent or AA but is slightly accelerated in its presence, (ii) acceleration is dependent on the presence of micelles irrespective of whether they are composed of mono-or polyunsaturated fatty acids, or of certain detergents, (iii) the acceleration is also observable when the N-terminal regulatory domain is removed, excluding that the open-closed conformational transition has any role in this effect, (iv), the same effect is seen when syntaxin 4 instead of syntaxin 3 is used, strongly arguing against a specific effect on syntaxin 3 as suggested in an earlier study (Darios and Davletov, 2006). I conclude that the observed acceleration of SNARE complex formation is a non-specific feature of *in-vitro* assembly that has no physiological significance. It is highly unlikely that free AA concentrations ever reach the CMC in cells and tissues: although I could not find data concerning free AA concentrations in cells and tissues, fatty acids are known to be not free in the cytoplasm but rather bound to proteins or else, inserted into membranes.

## Methods

### Constructs and protein purification

All bacterial expression constructs were derived from rat (*Rattus norvegicus*). The constructs for the SNARE proteins cysteine-free SNAP25 (1–206) and a variant thereof that carries a single cysteine at position 130 (SNAP-25^Cys130^), the cytoplasmic domains of syntaxin 3 (1–260), syntaxin 4 (1-273) and of synaptobrevin-2 (Syb2(1–96)), and the SNARE motifs of syntaxin 3 (183-261) and syntaxin 4 (192-279) were cloned into a pET28a vector that contains an *N*-terminal thrombin-cleavable His6 tag. All constructs were expressed in *Escherichia coli* BL21 DE3 cells. Proteins were purified by Ni^2+^-affinity chromatography, followed by ion-exchange chromatography essentially as previously described. SDS-PAGE was carried out as described by Laemmli. Nondenaturing gels were prepared and run in a manner identical to that of SDS-polyacrylamide gels except that SDS was omitted from all buffers as described (Fasshauer et al., 1999). All gels were stained by Coomassie-Blue.

### Fluorescence Spectroscopy

All measurements were carried out essentially as described(Fasshauer and Margittai, 2004; Wiederhold et al., 2010) in a Fluoromax 2 spectrometer equipped with autopolarizers (Jobin Yvon) in PBS buffer (20 mM phosphate buffer, pH 7.4, 150 mM NaCl, 1 mM DTT) at 25°C in 1-cm quartz cuvettes. Changes of anisotropy upon complex formation using proteins labeled with Oregon Green were measured at an excitation wavelength of 488 nm and an emission wavelength of 520 nm. The slit widths were set to 2– 4 nm, and the integration time was set at 1 s. The G factor was calculated according to G = I_HV_/I_HH_, where I is the fluorescence intensity, and the first subscript letter indicates the direction of the exciting light and the second subscript letter shows the direction of emitted light. The intensities of the vertically (V) and horizontally (H) polarized light emitted after excitation by vertically polarized light were measured. The anisotropy (r) was determined according to r=(I_VV_–G I_VH_)/(I_VV_ +2 G I_VH_).

### CD Spectroscopy

Measurements were performed essentially as described (Fasshauer et al., 2002; Wiederhold et al., 2010; Wiederhold and Fasshauer, 2009) using a Chirascan instrument (Applied Photophysics). For spectral measurements, different protein combinations at a concentration of about 5–10 μM in 20 mM Tris, pH 7.4, 100 mM NaCl (for details, see the appropriate figure legend) were incubated overnight. Hellma quartz cuvettes with a path length of 0.1 cm were used. The far-UV spectra were obtained using steps of 1 nm with a scan rate of 60 nm/min and an averaging time of 0.5–2 s. The measurements were carried out at 25 °C.

### Original data sets

The original data sets, including scan of the gels, fluorescence and circular dichroism data, can be found at https://doi.org/10.5281/zenodo.7610133.

## Abbreviations

(SNAREs): <u>N</u>-ethylmaleimide-<u>s</u>ensitive factor <u>a</u>ttachment <u>re</u>ceptors
(DDM): n-Dodecyl β-D-maltoside
(AA): arachidonic acid
(NGF): nerve growth factor
(CMC): critical micellar concentration

